# Development of an *in situ* cell-type specific proteome analysis method using antibody-mediated biotinylation

**DOI:** 10.1101/2023.06.13.544682

**Authors:** Taekyung Ryu, Seok-Young Kim, Thujitha Thuraisamy, Yura Jang, Chan Hyun Na

**Affiliations:** Department of Neurology, Johns Hopkins University School of Medicine, Baltimore, Maryland, USA; Neuroregeneration and Stem Cell Programs, Institute for Cell Engineering, Johns Hopkins University School of Medicine, Baltimore, Maryland, USA

**Keywords:** Cell-type-specific proteome analysis, mass spectrometry, brain, immunohistochemistry, biotin-tyramide, neuron, astrocytes, microglia

## Abstract

Since proteins are essential molecules exerting cellular functions, decoding proteome changes is the key to understanding the normal physiology and pathogenesis mechanism of various diseases. However, conventional proteomic studies are often conducted on tissue lumps, in which multiple cell types are entangled, presenting challenges in interpreting the biological dynamics among diverse cell types. While recent cell-specific proteome analysis techniques, like BONCAT, TurboID, and APEX, have emerged, their necessity for genetic modifications limits their usage. The alternative, laser capture microdissection (LCM), although it does not require genetic alterations, is labor-intensive, time-consuming, and requires specialized expertise, making it less suitable for large-scale studies. In this study, we develop the method for *in situ* <u>c</u>ell-type specific proteome analysis using <u>a</u>ntibody-mediated <u>b</u>iotinylation (iCAB), in which we combined immunohistochemistry (IHC) with the biotin-tyramide signal amplification approach. Poly-horseradish peroxidase (HRP) conjugated to the secondary antibody will be localized at a target cell type via a primary antibody specific to the target cell type and biotin-tyramide activated by HRP will biotinylate the nearby proteins. Therefore, the iCAB method can be applied to any tissues that can be used for IHC. As a proof-of-concept, we employed iCAB for mouse brain tissue enriching proteins for neuronal cell bodies, astrocytes, and microglia, followed by identifying the enriched proteins using 16-plex TMT-based proteomics. In total, we identified ∼8,400 and ∼6,200 proteins from enriched and non-enriched samples. Most proteins from the enriched samples showed differential expressions when we compared different cell type data, while there were no differentially expressed proteins from non-enriched samples. The cell type enrichment analysis with the increased proteins in respective cell types using Azimuth showed that neuronal cell bodies, astrocytes, and microglia data exhibited Glutamatergic Neuron, Astrocyte and Microglia/Perivascular Macrophage as the representative cell types, respectively. The proteome data of the enriched proteins showed similar subcellular distribution as non-enriched proteins, indicating that the iCAB-proteome is not biased toward any subcellular compartment. To our best knowledge, this study represents the first implementation of a cell-type-specific proteome analysis method using an antibody-mediated biotinylation approach. This development paves the way for the routine and widespread use of cell-type-specific proteome analysis. Ultimately, this could accelerate our understanding of biological and pathological phenomena.

## INTRODUCTION

Proteins are essential players and functional operatives within cells (1, 2). Therefore, the examination of proteome changes in the cell is essential to understanding normal physiology and pathogenesis processes of various diseases. Since the advent of the omics era, mass spectrometry-based proteomics has been instrumental in understanding proteome changes (3). While human or animal tissues are composed of multiple cell types intercalated with each other, the conventional proteomics method has been limited to reading the sum of their changes, compounding the interpretation of intricate proteome changes that occurred by multiple cell types (4).

To overcome these limitations, there have been many attempts for cell-type-specific proteome analysis. Alvarez-Castelao et al. reported a bioorthogonal noncanonical amino acid tagging (BONCAT)-based cell-type-specific proteome analysis approach, in which mutant type of methionyl-tRNA synthetase (MetRS*) that recognizes a noncanonical amino acid, azidonorleucine (ANL), is used to incorporate the noncanonical amino acid into the protein synthesis in a specific cell type that expresses MetRS*. The ANL-incorporated proteins could be biotinylated by click chemistry, retrieved by avidin beads and identified by mass spectrometry. They had MetRS* expressed in neuronal cells in a mouse brain and analyzed neuron-specific proteome changes when mice were exposed to an enriched environment, identifying 3,969 proteins using this approach (5). Hobson *et al*. developed a method to study the subcellular proteomics of dopaminergic neurons with cell-type specificity in the mouse brain using APEX2 proximity labeling and mass spectrometry (6). Shuster *et al*. reported in situ cell-surface proteome extraction by extracellular labeling (iPEEL) method, in which they had horse radish peroxidase (HRP) expressed on the cell surface in mouse organs and identified cell surface proteome. The biotinylated proteins could be enriched and identified in the same manner as described in the BONCAT method (7). Rayaprolu *et al*. reported a TurboID-based cell-type-specific proteome analysis method, in which they had TurboID expressed in a specific cell type of a mouse brain without a bait protein fused, and thereby they could label proteins in a specific cell type with biotin. The biotinylated proteins could be enriched and identified in the same way as described in BONCAT. Using the TurboID-based method, they performed the proteome analysis within distinct cell types, such as neurons and astrocytes. (8) In addition to the *in vivo* chemical tagging method, which labels proteins inside target cells or on cell surfaces, physical separation of target cells using laser capture microdissection (LCM) (9, 10) or fluorescence-activated cell sorting (FACS) (11) has been exploited too. However, the LCM method is tedious, time-consuming, and often needs histologist guidance. On the other hand, the FACS method needs the dissociation process of the target cells, which will lose a lot of *in vivo* proteome information in the process (12). As such, all these methods have the drawbacks of requiring the creation of genetically modified mouse lines or specialized instruments.

To overcome these limitations, we have developed the *in situ* <u>c</u>ell-type-specific proteome analysis method using <u>a</u>ntibody-mediated <u>b</u>iotinylation (iCAB), in which a target cell type can be bound by a primary antibody specific for the target cell and a secondary antibody conjugated with poly-HRP. Thereby, the cell-type-specifically localized HRP will activate nearby biotin-tyramide and, in turn, the activated biotin-tyramides biotinylate nearby proteins, achieving cell-type-specific biotinylation. The biotinylated proteins can be easily enriched using streptavidin-sepharose beads and identified by mass spectrometry analysis. Recently, this *in situ* antibody-based biotinylation method was also used for *in situ* interactome analysis, proving the broad applicability of this antibody-based biotinylation (13, 14). As a proof-of-concept experiment, we employed the iCAB method for neuronal cell bodies, astrocytes and microglia using anti-NeuN, anti-GFAP, and anti-IBA1 antibodies, respectively. To our knowledge, this is the first cell-type-specific proteome analysis using antibody-mediated biotinylation. This method is straightforward and user-friendly because any research laboratory where immunohistochemistry (IHC) is available can employ it. This method can be applied to any tissue section, including human tissues, with which IHC can be conducted. Therefore, iCAB will open a new avenue to the widespread use the cell-type-specific analysis, bringing the cell-type-specific proteome analysis to the level of routine proteome analysis.

## EXPERIMENTAL PROCEDURES

### Preparation of free-floating mouse brain sections

We purchased mouse brains immersed in 4% paraformaldehyde (PFA) (Rockland Immunochemical, Gilbertsville, PA, USA) and prepared mouse brain sections according to the previous report (15). Briefly, mouse brains were washed with 1× cold phosphate-buffered saline (PBS, Thermo Fisher Scientific, Cleveland, OH, USA), and immersed in 4% cold PFA in 0.1 M phosphate buffer (pH 7.4, Invitrogen, Carlsbad, CA, USA) for 2 days at 4 °C. After PFA immersion; the brains were incubated with 30% sucrose (Thermo Fisher Scientific) in PBS at 4°C, allowing them to sink to the bottom of the container completely. Subsequently, the mouse brains were sectioned coronally into 40 μm-thick slices at −18 to −20 °C using a microtome (Leica Microsystems, Wetzlar, Germany). Floating brain sections (fixed in 4% PFA) were stored in a cryoprotectant solution (ethylene glycol based; 30% ethylene glycol, 30% glycerol, and 10% 0.2 M sodium phosphate buffer pH 7.4, in distilled water (DW)) at −20 °C until use. All reagents related to cryoprotectant solution were purchased from Sigma-Aldrich (St. Louis, MO, USA)

### Cell-type-specific biotinylation using iCAB

Prepared floating brain sections were rinsed twice for 5 min with PBS and then were further washed with the wash buffer (0.2% Triton X-100 in PBS) for 10 min at room temperature on a low-speed rocker. The sections were incubated in a blocking buffer (0.2% of Triton X-100 and 5% of normal goat serum in PBS) for one hour at room temperature. Subsequently, primary antibodies were directly added to the floating sections in the blocking buffer in the following concentrations: rabbit anti-NeuN antibody (Invitrogen) for neuron at 1:500, rabbit anti-GFAP antibody (Invitrogen) for astrocyte at 1:100, and rabbit anti-IBA1 antibody (Wako) for microglia at 1:500. The sections treated with the primary antibody were incubated overnight at 4°C with low-speed rocker. We also prepared negative control samples that were treated the same way as other samples except those not treated with a primary antibody. Sections were then washed with the wash buffer thrice for 10 min each time and incubated with an poly-HRP-conjugated secondary antibody solution (SuperBoost™ Goat anti-Rabbit Poly HRP, IgG, Invitrogen) for 1 h at room temperature with gentle shaking. Sections were then washed with the wash buffer twice for 10 min each time, followed by washing with 0.1 M sodium borate buffer (pH 8.5) thrice for 10 min each time. For cell-type specific biotinylation, the sections were incubated with various concentrations of biotin-tyramide (Iris Biotech GmbH, Marktredwitz, Germany) in 0.1 M sodium borate (pH 8.5) for 30 min. Sections were then washed thrice for 10 min each time with wash buffer. Then, we incubated the section with the avidin-biotin complex (ABC) kit reagent (Vector Laboratories, Burlingame, CA) for 30 min to detect the cell-type-specific biotinylation of the mouse brain tissue. After washing with the wash buffer thrice for 10 min each time, sections were developed using Deep Space Black™ Chromogen Kit (Biocare Medical, Pacheco, CA, USA). The whole brain section images were acquired on the stereo microscope camera (AmScope, Irvine, CA, USA). The stained brain cell images were acquired on the optical microscope coupled with 16 MP USB 3.0 Color CMOS C-Mount Microscope Camera (AmScope).

### Enrichment, on-bead digestion and tandem mass tagging of cell-type-specifically biotinylated proteins

The biotinylated tissue sections were lysed with sonication in the lysis buffer consisting of 4% sodium dodecyl sulfate (SDS) and 1% sodium deoxycholate in 50 mM of ammonium bicarbonate (TEAB) buffer, followed by heating at 99 °C for an hour. Subsequently, lysate was centrifuged at 15,000 × g for five minutes, and the supernatant was reduced and alkylated using 10 mM tris(2-carboxyethyl)phosphine hydrochloride (TCEP, Sigma Aldrich) and 40 mM chloroacetamide (CAA, Sigma Aldrich) at room temperature for one hour.

To analyze the non-enriched samples, 10% of each lysate was acquired and subjected to methanol-chloroform precipitation to remove the SDS as follows. The 10% lysate was mixed with 4 volumes of 100% methanol and vortexed. Subsequently, 1 volume of chloroform was added, followed by vortexing. After adding 1 volume of DW, phase separation was induced by centrifugation at 12,000 × g for 10 min. The upper and lower layers were carefully removed, and the precipitated protein was washed with methanol. The washed protein was then digested with trypsin (sequencing grade modified trypsin; Promega) in a mixture of 100 mM ammonium bicarbonate (pH 8) and 2 M urea at the ratio of 50 to 1. The digestion was carried out at 37°C overnight. The obtained peptides for non-enrichment samples were desalted with C18 StageTips (3M Empore™, St. Paul, MN, USA).

To enrich the biotinylated proteins, the reduced and alkylated lysate was incubated with streptavidin-sepharose beads at room temperature for two hours with gentle rotation. Subsequently, the supernatant was removed from the beads, and the beads were washed five times with the first washing buffer (1% Triton X-100 and 0.2% SDS in DW). The beads were subsequently washed with the second washing buffer (50 mM ammonium bicarbonate in DW) twice. The biotinylated proteins bound to the beads were digested with sequencing-grade trypsin (Promega, Madison, WI, USA) in 100 mM ammonium bicarbonate (pH 8) and 2 M urea (at a ratio of 50:1) at 37 °C overnight. The digested peptides were collected and desalted with an SCX disk (3M Empore™). Both the digested peptides from non-enriched and enriched samples were labeled with 16-plex TMT according to the manufacturer’s instructions (Thermo Fisher Scientific). Briefly, the labeling reaction was carried out at room temperature for 1 hour, followed by quenching with 1/10 volume of 1 M Tris–HCl (pH 8.0). The obtained TMT-labeled peptides were pooled, vacuum dried using a SpeedVac (Thermo Fisher Scientific) and then stored at −80 °C until use.

### Prefractionation of peptides

The cell-type specific TMT-labeled peptides were reconstituted in 50 mM TEAB and underwent pre-fractionation using basic pH reversed-phase liquid chromatography (bRPLC), fractionating them in 96 fractions. Subsequently, these 96 fractions were concatenated into 24 fractions. For the basic pH reversed-phase liquid chromatography fractionation, an Agilent 1260 offline LC system (Agilent Technologies) was employed, which consisted of a binary pump, UV detector, an autosampler, and an automatic fraction collector. Briefly, the dried peptide samples were reconstituted in solvent A (10 mM TEAB in water, pH 8.5) and loaded onto an Agilent 300 Extend-C18 column (5 μm, 4.6 mm × 25 cm; Agilent Technologies). The peptides were then resolved using an increasing gradient of solvent B (10 mM TEAB in 90% acetonitrile (ACN), pH 8.5) at a flow rate of 0.3 mL/min, with a total run time of 150 min. Finally, the 24 concatenated samples were vacuum-dried using a SpeedVac and stored at −80°C until further use.

### Mass spectrometry analysis

Liquid chromatography with tandem mass spectrometry (LC-MS/MS) analysis was conducted as described previously with minor modifications (16, 17). The Orbitrap Fusion Lumos Tribrid Mass Spectrometer (Thermo Fisher Scientific) coupled with an Ultimate 3000 RSLCnano nanoflow liquid chromatography system (Thermo Fisher Scientific) was used to analyze fractionated peptides. The peptides from each fraction were reconstituted in 0.5% formic acid (FA) and loaded onto a trap column (Acclaim PepMap 100, LC C18, 5 μm, 100 μm × 2 cm, nanoViper) at a flow rate of 8 μL/min. The peptides were resolved at a flow rate of 0.3 μL/min using an increasing gradient of solvent B (0.1% FA in 95% ACN) on an analytical column (Easy-Spray PepMap RSLC C18, 2 μm, 75 μm × 50 cm) equipped with an EASY-Spray ion source that was operated at a voltage of about 2.4 kV. The total run time was 120 min. Mass spectrometry (MS) analysis was performed in data-dependent acquisition (DDA) mode with a full scan in the mass-to-charge ratio (*m/z*) range of 300 to 1800 in the “Top Speed” mode with 3 s per cycle. MS1 and MS2 scans were acquired for the precursor and the peptide fragmentation ions, respectively. MS1 scans were measured at a resolution of 120,000 at an *m/z* of 200, while MS2 scans were acquired by fragmenting precursor ions using the higher-energy collisional dissociation (HCD) method, which was set to 35% of collision energy and detected at a mass resolution of 50,000 at an *m/z* of 200. The automatic gain control targets were set to one million ions for MS1 and 0.05 million ions for MS2. The maximum ion injection time was set to 50 ms for MS1 and 100 ms for MS2. The precursor isolation window was set to 1.6 *m/z* with 0.4 *m/z* of offset. Dynamic exclusion was set to 30 s, and singly charged ions were rejected. Internal calibration was carried out using the lock mass option (*m/z* 445.12002) from ambient air.

### Database searches for cell-type specific proteomes

Database searches were conducted as described previously with minor modifications (16, 17). The MS/MS data obtained from LC-MS/MS analyses were used to search against the UniProt mouse protein database (UP000000589, included both Swiss-Prot and TrEMBL and released in January 2019 with 55,435 entries), which contained protein entries with common contaminants (115 entries), using MSfragger 3.4 (18) search algorithms through Thermo Proteome Discoverer software package (version 2.4.1.15, Thermo Scientific). The top ten peaks in each window of 100 *m/z* were selected for database search during MS2 preprocessing. The following parameters were used for the database search. Trypsin was set as the protease, allowing for a maximum of two missed cleavages. Carbamidomethylation of cysteine (+57.02146 Da) and TMTpro tags (+304.20715 Da) on lysine and peptide N termini were set as fixed modifications, while oxidation (+15.99492 Da) of methionine and biotinylation (+361.14600 Da) of tyrosine were set as a variable modification. The minimum peptide length was set to six amino acids. The precursor mass (MS1) and fragment mass (MS2) tolerances were set to 10 ppm and 20 ppm, respectively. Peptides and proteins were filtered at a 1% false discovery rate (FDR) using the percolator node and protein FDR validator node, respectively. The following parameters were used for the protein quantification. The most confident centroid option was used for the integration mode, while the reporter ion tolerance was set to 20 ppm. The MS order was set to MS2, and the activation type was set to HCD. Both unique and razor peptides were used for peptide quantification, while protein groups were considered for peptide uniqueness. The coisolation threshold was set to 50%. Reporter ion abundance was computed based on signal-to-noise ratios, and the missing intensity values were replaced with the minimum value. The average reporter signal-to-noise threshold was set to 10. The quantification value corrections for isobaric tags and data normalization were disabled. Protein grouping was performed with a strict parsimony principle to generate the final protein groups. All proteins sharing the same set or subset of identified peptides were grouped, while protein groups with no unique peptides were filtered out. Proteome Discoverer iterated through all spectra and selected PSMs with the highest number of unambiguous and unique peptides, and then final protein groups were generated. The Proteome Discoverer summed all the reporter ion abundances of PSMs for the corresponding proteins (19).

### Bioinformatics and statistical analysis

The statistical analysis of the mass spectrometry data was performed with the Perseus software (version 1.6.0.7) (20). For normalization, the reporter ion intensity values were divided by the median values of each protein, and the relative abundance values for each sample were subtracted by the median values of each sample after the log2 transformation. The *p* values between the comparison groups were calculated using the student’s two-sample *t*-test. Proteins with *q* < 0.05 were considered differentially expressed. The fold change was calculated by dividing the mean value of one group by that of another group. The *q-*values for the volcano plot were calculated by significance analysis of microarray (SAM) and a permutation-based FDR estimation (21). We used the Azimuth database embedded in Enrichr to discover the closes cell type for the input list of genes (22). Azimuth is an annotated reference dataset constructed based on single-cell RNA-seq data (23). The development of Azimuth is led by the New York Genome Center Mapping Component as part of the NIH Human Biomolecular Atlas Project (HuBMAP) (24). We used the top 300 upregulated proteins with the *q*-value < 0.05 from a specific cell type for the Azimuth cell-type analysis. Principal Component Analysis (PCA) and heatmap analyses were conducted using the MetaboAnalyst tool (25). Gene Ontology Cellular Component (GO-CC) analysis embedded in DAVID bioinformatics resources (version 6.8) was used to examine the distribution pattern of the input proteins in the cell (26). For the GO-CC analysis, we used all the proteins identified by 3 or more peptides.

## RESULTS

The primary objective of this research is to develop a straightforward and user-friendly method for cell-type-specific proteomics analysis for any tissue sections combining IHC with the biotin-tyramide signal amplification method. As a proof of concept, we employed iCAB to a mouse brain, in which various cell types are entangled in a complex structure. For example, to conduct cell-type-specific proteome analysis for astrocytes, we used a rabbit anti-GFAP antibody followed by an anti-rabbit antibody conjugated with poly-HRP. Then, HRP localized at astrocytes will make biotin-tyramide locally activated and, subsequently, the locally activated biotin-tyramide will biotinylate nearby proteins on tyrosine residues, resulting in the biotinylation of the astrocyte proteins. The level of biotinylation can be confirmed with chromogen. Either stained sections by chromogen or non-stained sections can be used for the enrichment of biotinylated proteins using streptavidin beads, followed by the identification of biotinylated proteins using mass spectrometry (Figure 1).

**Figure 1.**
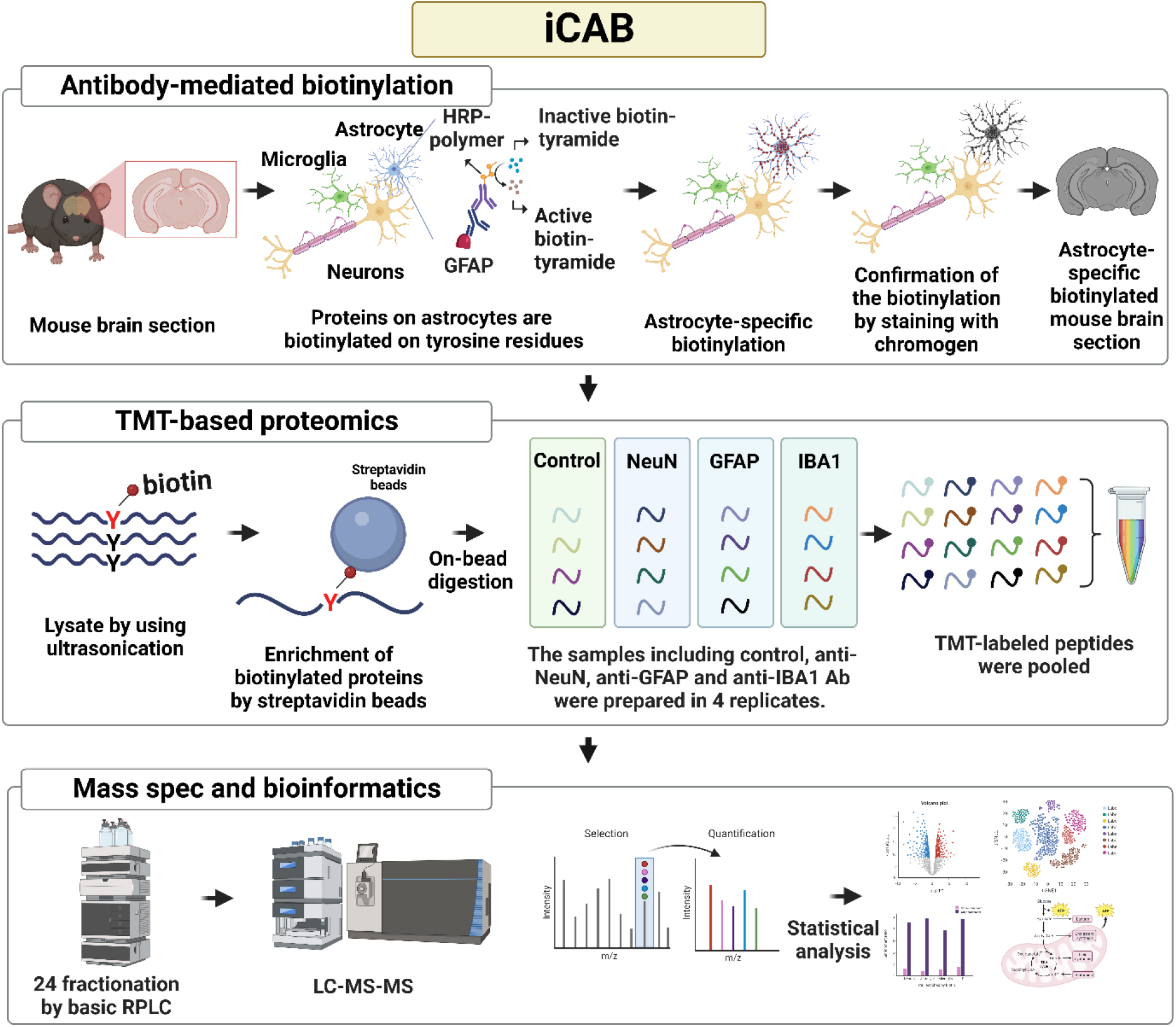
Experimental strategy to identify neuronal cell body-, astrocyte- and microglia-specific proteomes using iCAB method. The iCAB method incorporates antibody-mediated biotinylation and quantitative proteome analysis, facilitating the identification of cell-type-specific proteins. The process of antibody-mediated biotinylation involves targeting a specific cell type using a primary antibody specific to the target cell type and a secondary antibody conjugated with poly-HRP. Biotinylation is accomplished by activating biotin-tyramide using poly-HRP in the presence of hydrogen peroxide. To confirm cell-type-specific biotinylation, we used an ABC kit containing streptavidin and biotinylated HRP, followed by color development with Deep Space Black chromogen. For quantitative TMT-based proteome analysis, we enriched biotinylated proteins using streptavidin-sepharose beads and conducted on-bead digestion. The resulting peptides were labeled with 16-plex TMT, prefractionated by bRPLC, and analyzed by LC-MS/MS.

### Biotinylation of mouse brain sections using iCAB

To conduct the cell-type-specific biotinylation, we prepared PFA-fixed wild-type mouse brain sections with 40 μm thickness. The PFA-fixed brain tissue sections were incubated with anti-NeuN, anti-GFAP, and anti-IBA1 antibodies to target neuronal cell bodies, astrocytes, and microglia, respectively. Subsequently, the primary antibodies were detected with the secondary antibody conjugated with poly-HRP. To investigate the optimal antibody concentration, we tested various antibody concentrations. Anti-NeuN, anti-GFAP, and anti-IBA1 antibodies showed the best staining contrast at 1:500, 1:500, and 1:100 dilution ratios, respectively (Supplemental Figure S1). We decided to use these antibody concentrations for the follow-up experiments. The tissue sections bound by the primary and secondary antibodies were biotinylated cell-type-specifically using biotin-tyramide and hydrogen peroxide. To ascertain the most efficacious concentration of biotin-tyramide, we conducted biotinylation procedures utilizing varying concentrations of biotin-tyramide; 0, 0.3, 3, 30, and 150 μM. The level of biotinylation was visualized using avidin/biotin-HRP complex (ABC) kit in the presence of hydrogen peroxidase and subsequently deep space black chromogen. Before staining with ABC kit, we depleted the residual HRP from the secondary antibody by treating the tissue sections with 3% hydrogen peroxide to avoid the possible unwanted staining by the residual HRP from the secondary antibody. As expected, the brain tissue section without biotin-tyramide did not show any staining, while other brain tissue sections showed biotin-tyramide concentration-dependent coloring, supporting that the staining was biotinylation-dependent (Supplemental Figure S2). Although 150 μM showed the darkest staining, 3 μM showed the best staining contrast for all three cell types, and we decided to use 3 μM for the biotin-tyramide staining for all three cell types. For the tissue sections stained with the optimal conditions, we further scrutinized the staining pattern of the target cells on various regions, such as the frontoparietal cortex, hippocampus, and thalamus. As expected, the iCAB staining using anti-NeuN antibody exhibited round shapes suggesting the staining of the neuronal cell bodies, the iCAB staining using anti-GFAP antibody exhibited star shapes with a lot of spikes suggesting the staining of astrocytes, and the iCAB staining pattern using anti-IBA1 antibody exhibited smaller star shapes with a lot of branches suggesting the staining of microglia (Figure 2). These results demonstrated that our cell-type-specific biotinylation method was successfully developed and optimized.

**Figure 2.**
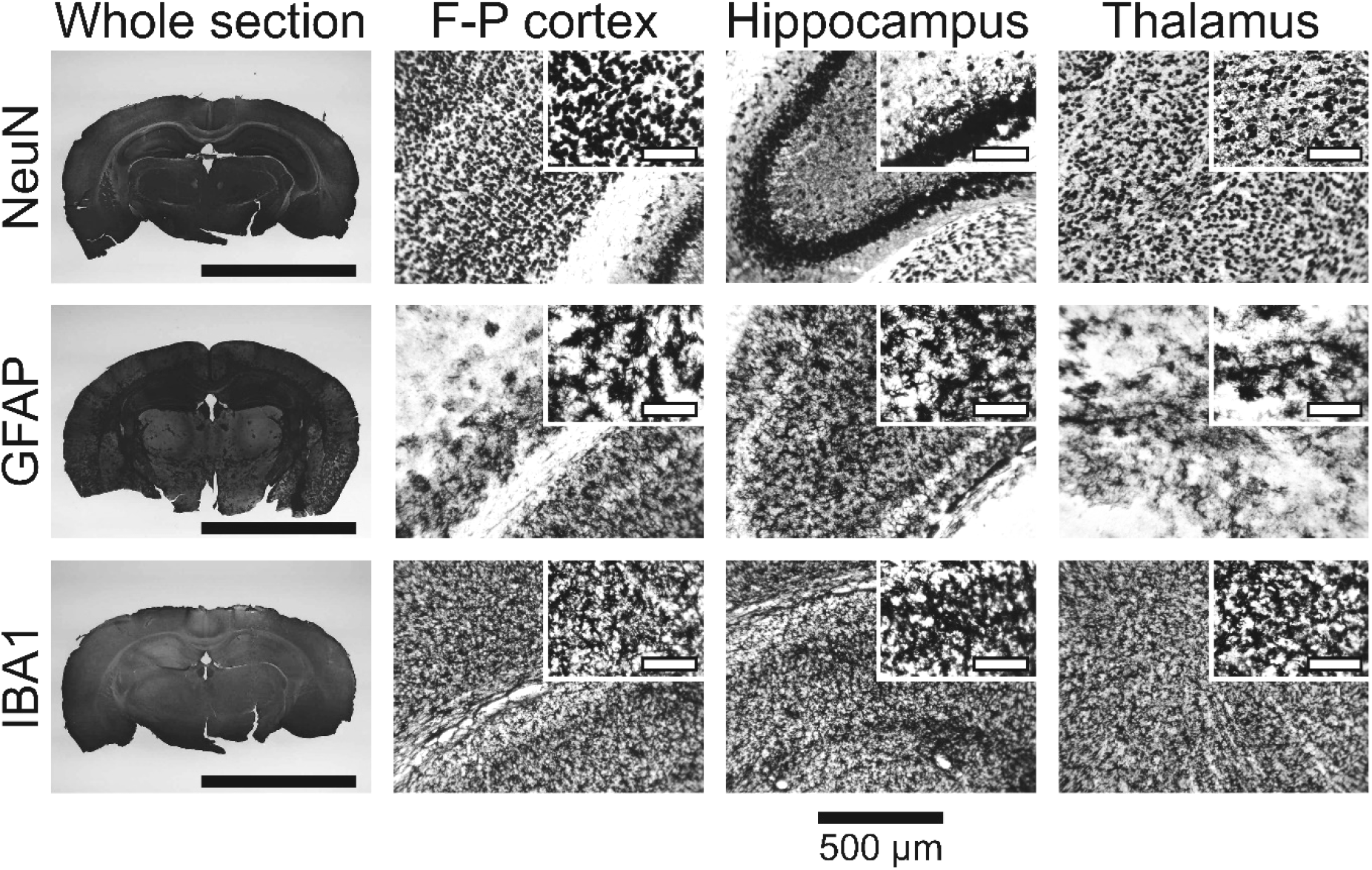
Chromogen-stained images of the biotinylated neuronal cell bodies, astrocytes and microglia in the frontoparietal (F-P) cortex, hippocampus and thalamus region. Neuronal cell bodies, astrocytes, and microglia proteins were biotinylated by anti-NeuN antibody (NeuN), anti-GFAP antibody (GFAP), and anti-IBA1 antibody (IBA1), respectively, using the iCAB method. The biotinylated proteins from each cell-type were detected by ABC kit composed of streptavidin and biotinylated HRP. The dark color on each cell body was developed by Deep Space Black chromogen. The insets in F-P cortex, hippocampus and thalamus regions are for magnified images, where the scale bar length is 100 μm. The scale bar length at the bottom of the whole section images is 5 mm.

### Mass spectrometry analysis for the identification of the cell-type-specifically biotinylated proteins

To identify the cell-type-specifically biotinylated proteins, we prepared 40 tissue sections with 40 μm thickness per replicate for anti-NeuN antibody (NeuN group), anti-GFAP antibody (GFAP group), anti-IBA1 antibody (IBA1 group), and no primary antibody (the control group). We included the negative control group, which was not treated with the primary antibody to evaluate non-specific binding to streptavidin-sepharose beads. The experiment was conducted in 4 replicates per group. The IHC and biotinylation were conducted with the best condition optimized. We stained one of the 4 replicates with the chromogen to confirm the proper biotinylation. All the tissue sections were lysed by sonication followed by heating to de-crosslink PFA linkages on proteins. The tissue section lysates were incubated with streptavidin-sepharose beads to enrich biotinylated proteins, followed by washing to remove non-specific binding proteins. Then, the bound proteins were released by on-bead digestion. The resulting digested peptides were labeled by 16-plex TMT to multiplex 4 replicates of the NeuN, GFAP, IBA1, and control groups. When the 16 samples were labeled by TMT, we randomized the order of the labeling to avoid any potential experimental bias. The labeled samples were pooled and prefractionated by bRPLC into 24 fractions for in-depth proteome analysis. To evaluate the enrichment efficiency, we saved a portion of each sample before enriching the biotinylated proteins and the samples were subjected to labeling with 16-plex TMT and bRPLC fractionation. Both non-enriched and enriched samples were analyzed by LC-MS/MS. From these LC-MS/MS analyses, we acquired 908,222 and 998,024 MS/MS spectra from non-enriched and enriched samples, respectively. After database search analysis, we could identify 43,059 peptides and 6,213 proteins from non-enriched samples and 67,956 peptides and 8,430 proteins from the enriched samples. The numbers of quantified proteins were 5,966 and 8,214 for non-enriched and enriched samples, respectively. We then visualized the differences between samples on PCA plots. While the non-enriched samples did not show any clustering on 2D PCA plane, the enriched samples showed clear clustering for each cell type group on 2D PCA plane (Supplementary Figure S3). The heap map analysis also showed that the enriched samples showed clear clustering for each group compared to the non-enriched samples (Supplemental Figure S4). In this TMT-based proteome analysis of the enriched cell types, one out of four samples per each group was stained by chromogen to ascertain the biotinylation level. Interestingly, the chromogen-stained sample showed no difference from three other non-stained samples, demonstrating that either stained or non-stained tissue sections can be used for the downstream proteome analysis without significant differences. These results demonstrate successful distinctive enrichments for 4 different groups.

### Statistical analysis of the quantified proteins

To identify the differentially expressed proteins in each enrichment group, we conducted statistical analysis for both non-enriched and enriched samples to estimate the enrichment efficiency, as described in the method section. The non-enriched samples showed no differentially expressed proteins in any comparisons between groups. On the other hand, almost all proteins for the enriched samples exhibited differential expression (Figure 3). In the comparison of the GFAP group to IBA1 group, we identified 3,798 and 3,907 upregulated proteins in the GFAP and IBA1 groups, respectively. In the comparison of the GFAP group to the NeuN group, we identified 3,946 and 3,863 upregulated proteins in the GFAP and NeuN groups, respectively. In the comparison of the NeuN group to the IBA1 group, we identified 2,898 and 4,771 upregulated proteins in the NeuN and IBA1 groups, respectively. Notably, the cell-type marker proteins such as GFAP, AIF1 (IBA1) and RBFOX3 (NeuN) exhibited differential expressions (Figure 3). However, when the NeuN group is compared to the GFAP group, the Z score for RBFOX3 (NeuN) was 2, while some proteins showed a Z score > 8. This marginal enrichment of RBFOX3 can potentially be explained by the close entanglement of astrocytes with neuronal cell bodies (27). We observed similar findings when one cell type was compared to the control or two other cell types. (Supplementary Figures S5 and S6). These data support that the iCAB method efficiently enriched the biotinylated proteins cell-type-specifically.

**Figure 3.**
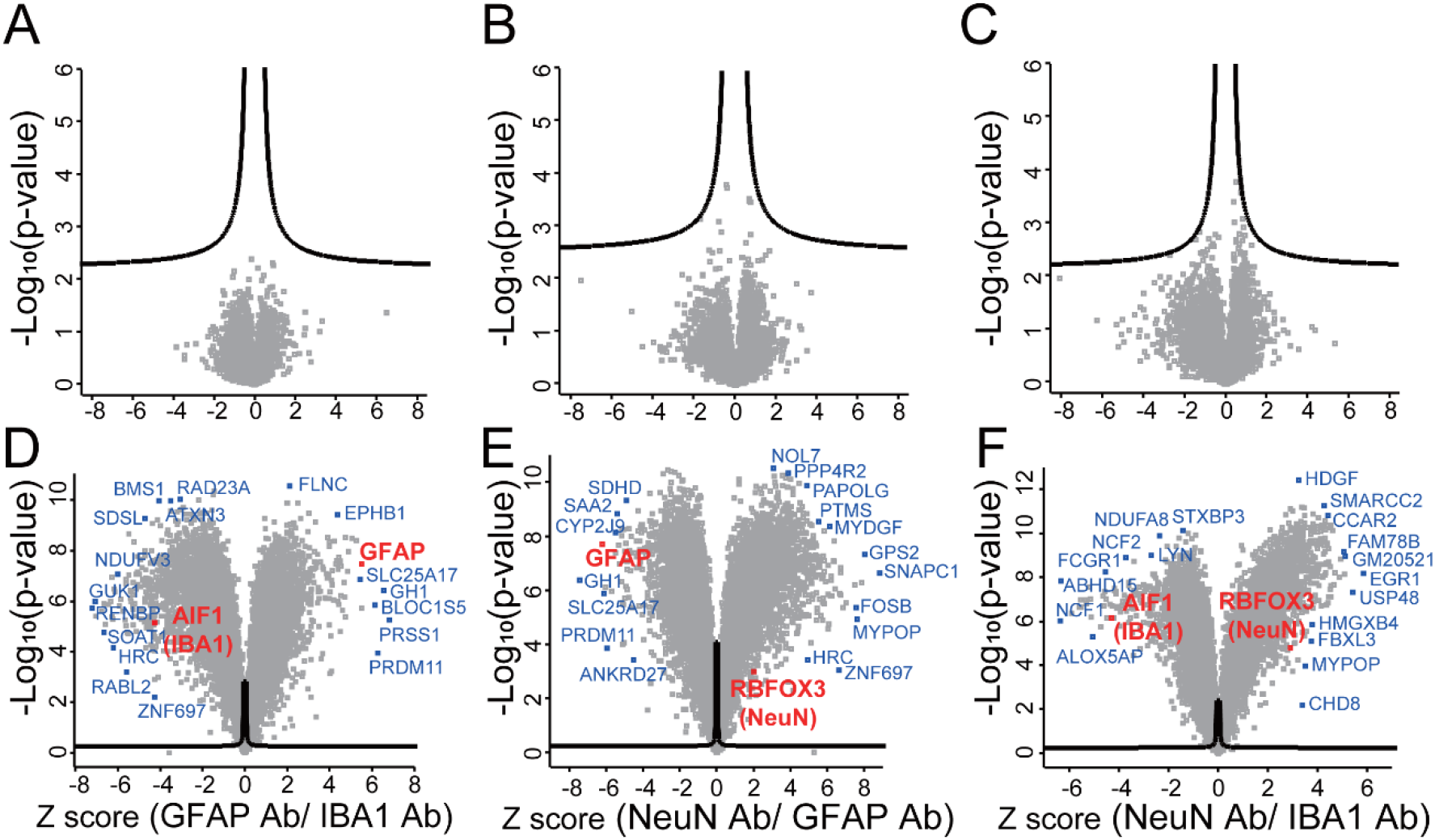
Volcano plot analysis results of the iCAB proteomics data. The three volcano plots on the top (A-C) are of iCAB proteome analysis results for non-enriched samples. The three volcano plots on the bottom (D-F) are of iCAB proteome analysis results for enriched samples. The iCAB proteome data of astrocytes were compared to that of microglia (A and D). The iCAB proteome data of neuronal cell bodies were compared to that of astrocytes (B and E). The iCAB proteome data of neuronal cell bodies were compared to that of microglia (C and F). The proteins outside the curved lines have *q*-values < 0.05.

### Cell-type enrichment analysis

We next investigated whether the upregulated proteins in each cell type showed any correlation with the respective cell type. We first selected the top 300 proteins that show the most upregulation in each cell type compared to two other cell types among the proteins that show *q*-value < 0.05. Subsequently, the selected top 300 proteins were fed to the Azimuth annotation reference dataset embedded in Enrichr. In alignment with our expectations, the top 300 proteins upregulated in neuronal cell bodies, astrocytes and microglia showed the biggest enrichment toward Glutamatergic neurons, Astrocyte1, and Microglia/Perivascular Macrophage, respectively (Figure 4). This data also demonstrates that the iCAB successfully enriched proteins from respective cell types.

**Figure 4.**
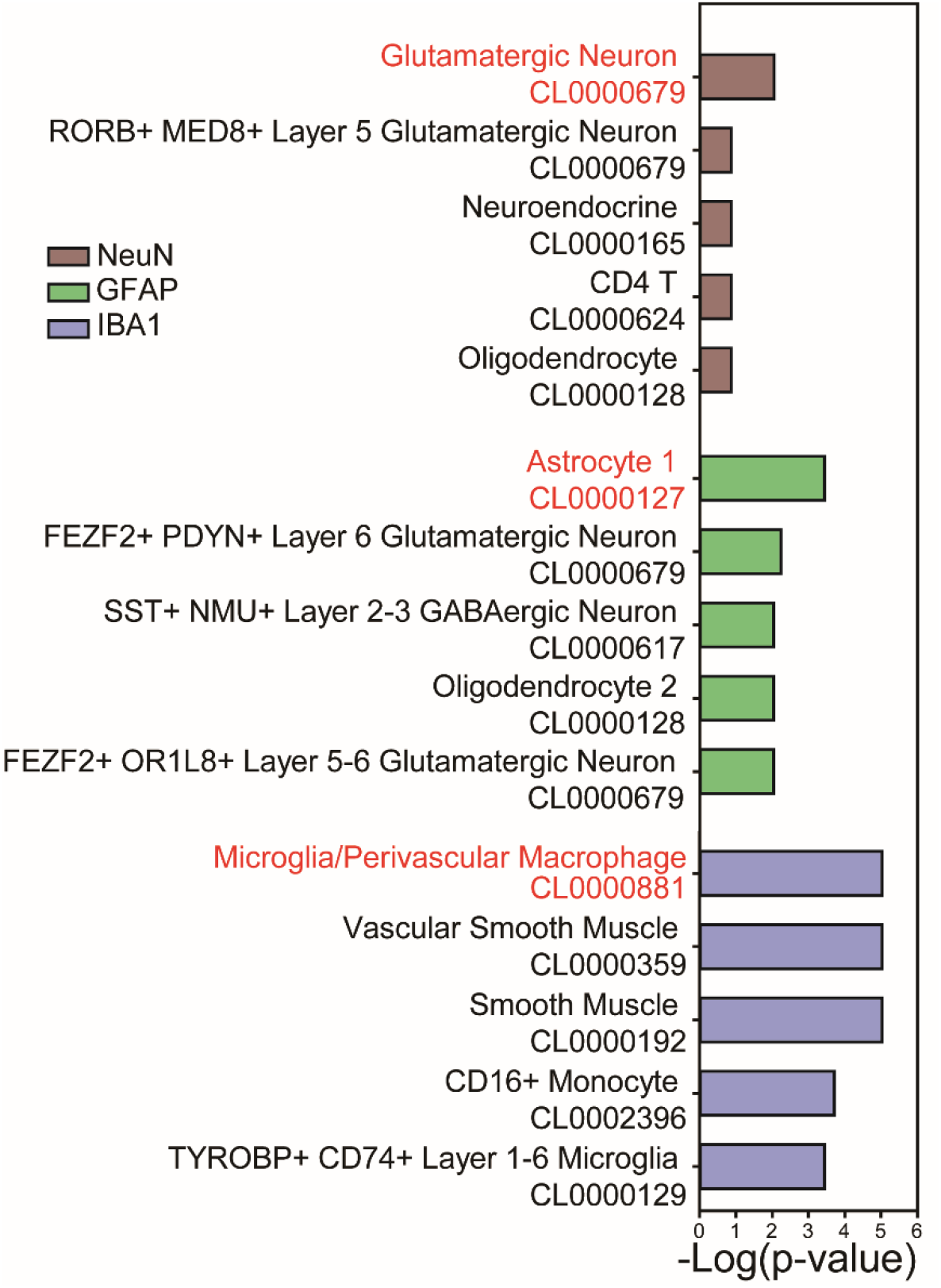
Cell-type-enrichment analysis using the Azimuth Cell-Type database. To identify enriched cell types for the upregulated proteins of the iCAB proteomics data, a cell-type-enrichment analysis was conducted using the Azimuth Cell-Type database. Proteins upregulated in a specific cell type were selected by comparing iCAB proteome data of a cell type to that of the other two cell types. Subsequently, the top 300 upregulated proteins were selected based on their Z scores among the proteins with *q*-values < 0.05.

### Distribution of subcellular localization of the cell-type specific proteome

We next investigated whether the iCAB method effectively biotinylated proteins evenly across the entire cellular compartments without any bias toward a certain compartment. For this, we performed Gene Ontology analysis using cellular compartment terms for both enriched and non-enriched proteomes. As shown in Figure 5, both enriched and non-enriched proteomes showed quite similar distribution patterns across various subcellular compartments, including the nucleus, cytosol, plasma membrane, mitochondria, and endomembrane system. This result suggests that the iCAB method successfully achieved biotinylation evenly across diverse subcellular compartments without any noticeable bias towards a specific organelle or compartment.

**Figure 5.**
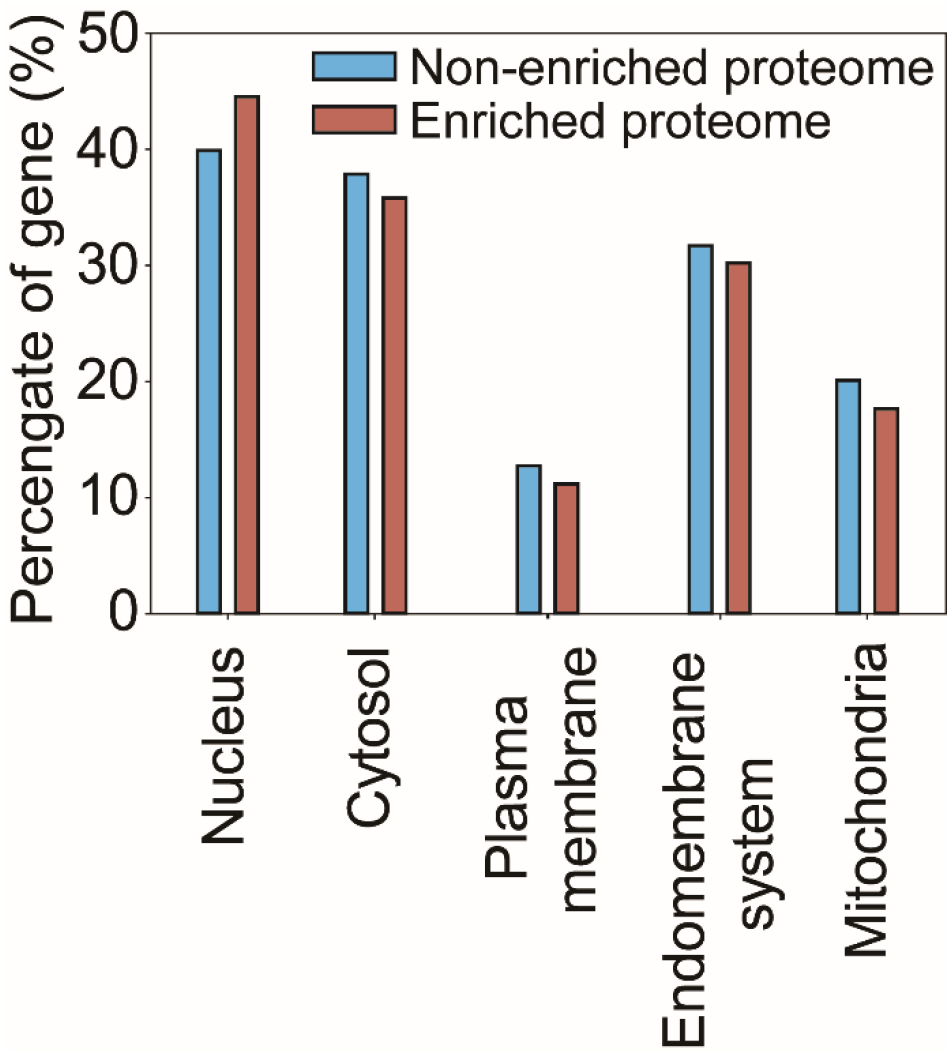
The subcellular localization of the identified proteins determined by Gene Ontology Cellular Compartment (GOCC) annotations. The percentage of gene represents the ratio of identified proteins that belong to the corresponding subcellular compartment. GO analysis was performed using all identified proteins identified three or more peptides.

## DISCUSSION

In this study, we aimed to devise a straightforward and user-friendly method to carry out cell-type-specific proteomics analysis for human or animal tissue sections. The iCAB method developed herein effectively fulfills this objective, as validated by the proof-of-concept testing using mouse brain tissue sections. By applying various antibodies targeting different cell types, followed by HRP-linked secondary antibodies, we could successfully localize HRP at the desired cell type, inducing biotinylation on tyrosine residues cell-type-specifically. This method was optimized and substantiated by visualizing the biotinylation levels with the ABC kit, which stained the biotinylated proteins. Crucially, the iCAB method could distinguish between different cell types, as seen in the differential biotinylation in neuronal cell bodies, astrocytes, and microglia. The use of mass spectrometry further helped to identify these biotinylated proteins, providing a quantifiable measure of the protein distribution cell type specifically. The successful implementation of the iCAB method led to notable disparities in protein abundance distinct to each cell type. Especially, the sample stained with chromogen did not show noticeable differences from non-stained samples in protein abundance, and this demonstrates that the iCAB users can validate the biotinylation level by chromogen staining whenever required without concerning the compromised proteome quality.

In this study, we identified ∼8,400 proteins from the iCAB-enriched samples. Since this number is quite close to the number of proteins identified in common brain proteome analysis without cell-type specific enrichment (28), this large number of protein identification also bolsters the strength of the iCAB method. Since the brain sections we used were fixed with PFA, potentially, the incomplete decrosslinking affected the number of identifications. This suggests that we will have more room for improvement in the number of proteins that can be identified.

Statistical analysis revealed differential expressions for almost all proteins in enriched samples, underscoring the capability of iCAB to effectively isolate cell-type-specific proteomes. However, in the case of RBFOX3 (NeuN), we observed a marginal enrichment when comparing neuronal cell body proteome to astrocyte proteome. This could potentially be explained by the close interweaving of astrocytes and neuronal cell bodies, which may affect the specificity of the enrichment. This could be improved by further optimization of the biotinylation resolution. When too many biotin-tyramides are activated, the activated biotin-tyramide can go over the boundary of each cell type and can biotinylate proteins in the intertwined neighboring cells. If we reduce the level of the biotin-tyramide activation, this could potentially be controlled.

Our Azimuth cell-type analysis further validated the efficacy of the iCAB method. Proteins highly upregulated in each cell type correlated strongly with the expression profiles of respective cell types in the Azimuth analysis. This data supports that the method can successfully enrich proteins from specific cell types, further strengthening its utility in cell-type-specific proteomic analysis. Moreover, Gene Ontology analysis of both enriched and non-enriched proteomes revealed a similar distribution pattern of proteins across various subcellular compartments. This suggests that iCAB efficiently biotinylated proteins across the entire cell without a noticeable bias towards a specific organelle, underlining the comprehensiveness of this method. As demonstrated so far, the iCAB method is versatile and can be applied to different cell types and subcellular organelles in any organ regardless of species, including human tissues, given the availability of a primary antibody that specifically targets the desired antigen.

More importantly, since iCAB has been developed with a minor modification of IHC by adopting biotin-tyramide signal amplification, almost all biomedical research labs can adapt their IHC to this iCAB without much optimization effort. Once the proteins on a target cell type are biotinylated, they can be easily enriched and identified by common proteomics facilities in most research institutes. This flexibility and user-friendliness allow its application across diverse disease types, enabling invaluable insights into cell-cell interactions and their function involved in disease pathology. iCAB method opens up new avenues for biomedical research by bringing down the cell-type-specific proteome analysis to the level of routine proteome analysis.

## Author contributions

C.H.N. conceived the iCAB method. C.H.N, T.R., S.Y.K., and T.T. designed the experiments. T.R., S.Y.K., T.T, and Y.J. performed the experiments and analyzed the data. T.R., S.Y.K., and C.H.N wrote the manuscript. C.H.N. supervised the research.

## Funding and additional information

This study was supported by the National Institutes of Health grant (R01NS123456 to C.H.N.). We acknowledge the National Institutes of Health shared instrumentation grant (S10OD021844).

---

**Supplementary Figure S1.**
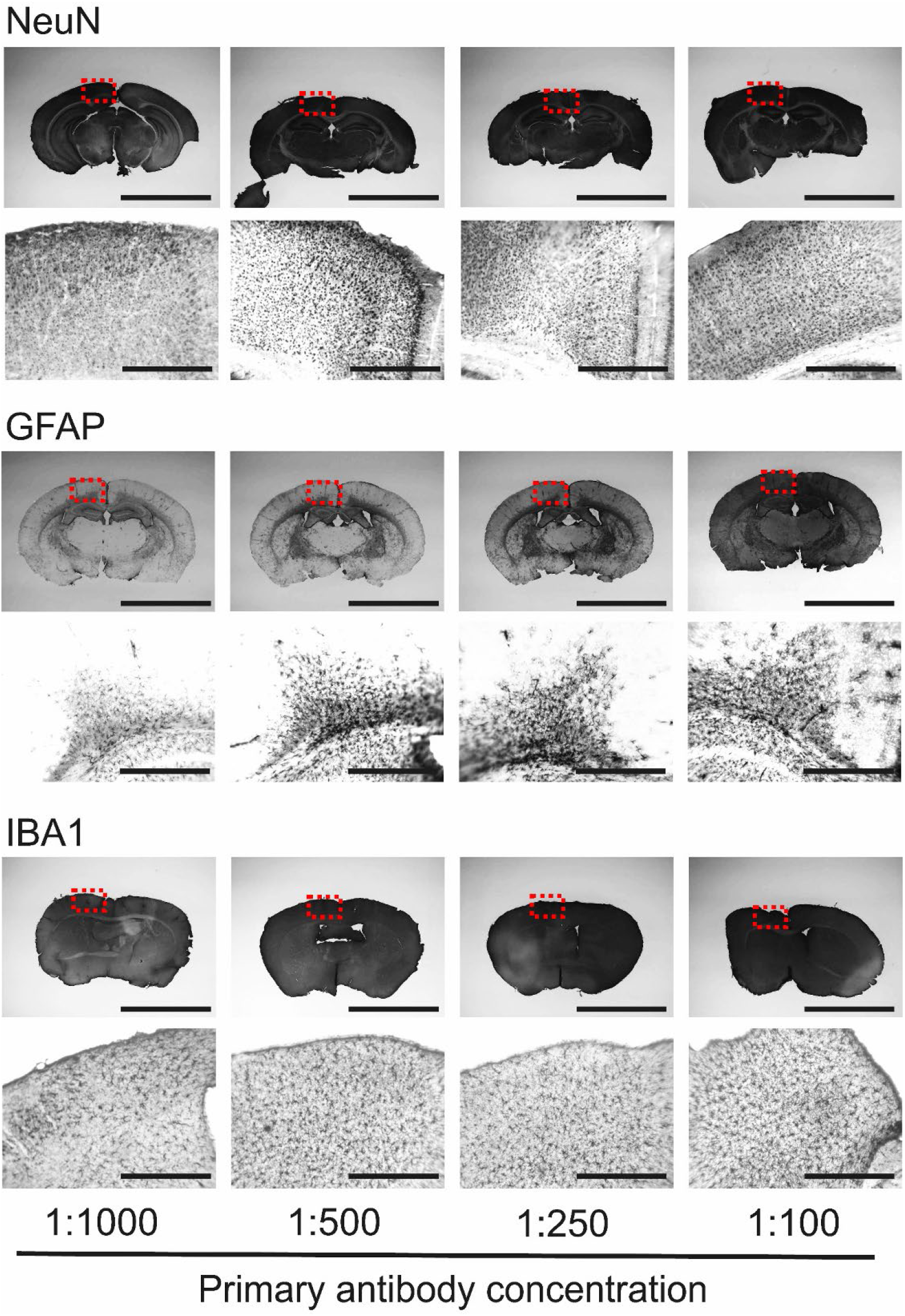
Optimization of the iCAB method for the primary antibody concentrations. Different concentrations of anti-NeuN antibody (NeuN), anti-GFAP antibody (GFAP), and anti-IBA1 antibody (IBA1) were tested for iCAB. The images on the top are of whole brain sections and the images on the bottom are zoom-ins of the red dotted squares on the top. The lengths of the scale bars for the whole sections (top) and the F-P cortices (bottom) are 5 mm and 500 μm, respectively.

**Supplementary Figure S2.**
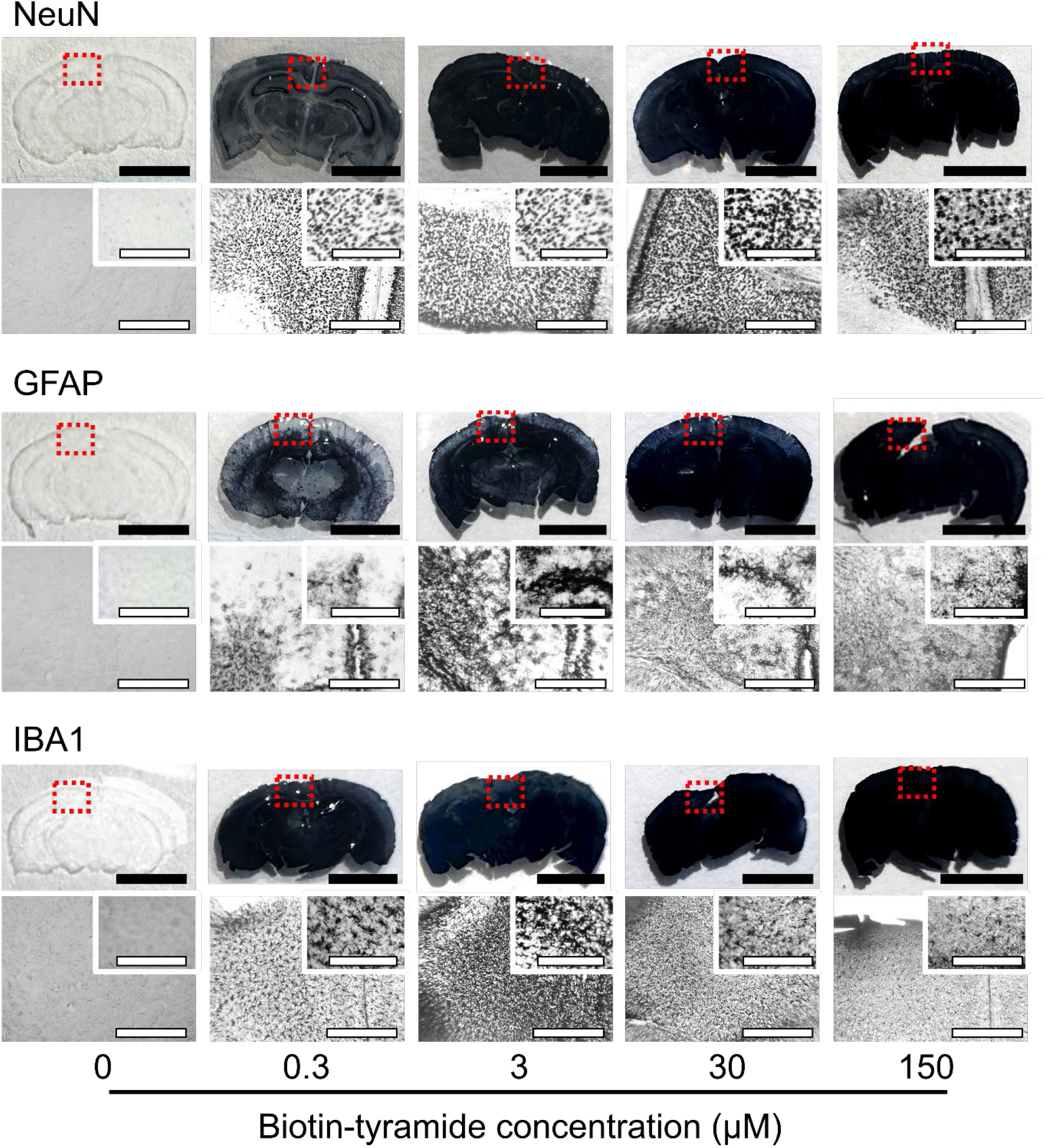
Optimization of the iCAB method for the biotin-tyramide concentrations. Five different concentrations of biotin-tyramide for anti-NeuN antibody (NeuN), anti-GFAP antibody (GFAP), and anti-IBA1 antibody (IBA1) were evaluated to find the optimal biotin-tyramide concentrations for them. The images on the top are of whole brain sections and the images on the bottom are zoom-ins of the red dotted squares on the top. The lengths of the scale bars for the whole sections (top), P-F cortices (bottom) and the insets of P-F cortices (the top-right corner of the images on the bottom) are 5 mm, 500 μm, and 100 μm, respectively.

**Supplementary Figure S3.**
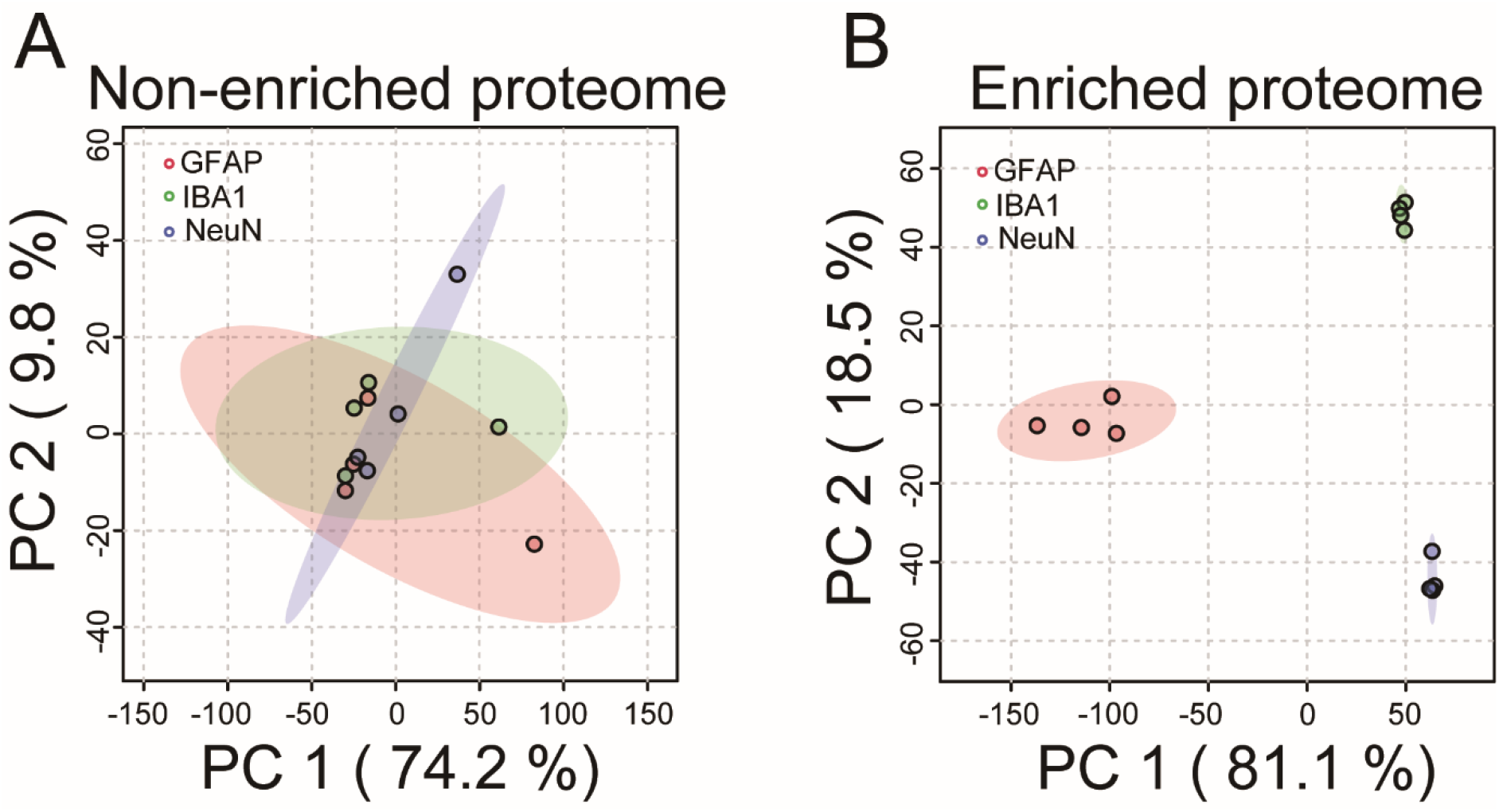
Principal component analysis (PCA) of the iCAB proteome data. The PCA analyses for the non-enriched (A) and enriched (B) proteome data were conducted. GFAP, IBA1 and NeuN represent the proteome data for astrocytes (anti-GFAP antibody), microglia (anti-IBA1 antibody), and neuronal cell body (anti-NeuN antibody).

**Supplementary Figure S4.**
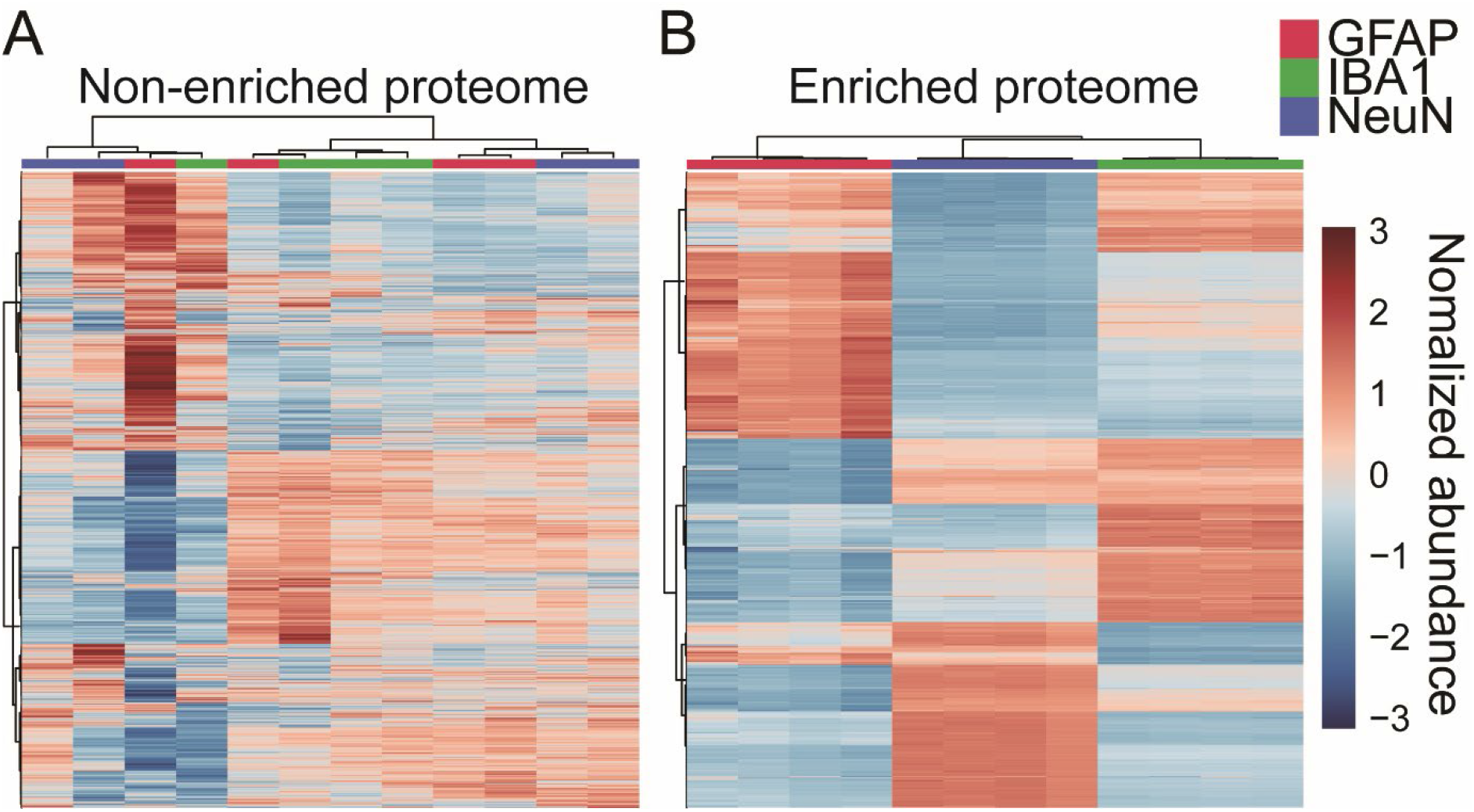
Hierarchical-clustered heatmap analysis. Hierarchical-clustered heatmaps for non-enriched (A) and enriched (B) samples were generated. GFAP, IBA1 and NeuN represent the proteome data for astrocytes (anti-GFAP antibody), microglia (anti-IBA1 antibody), and neuronal cell body (anti-NeuN antibody).

**Supplementary Figure S5.**
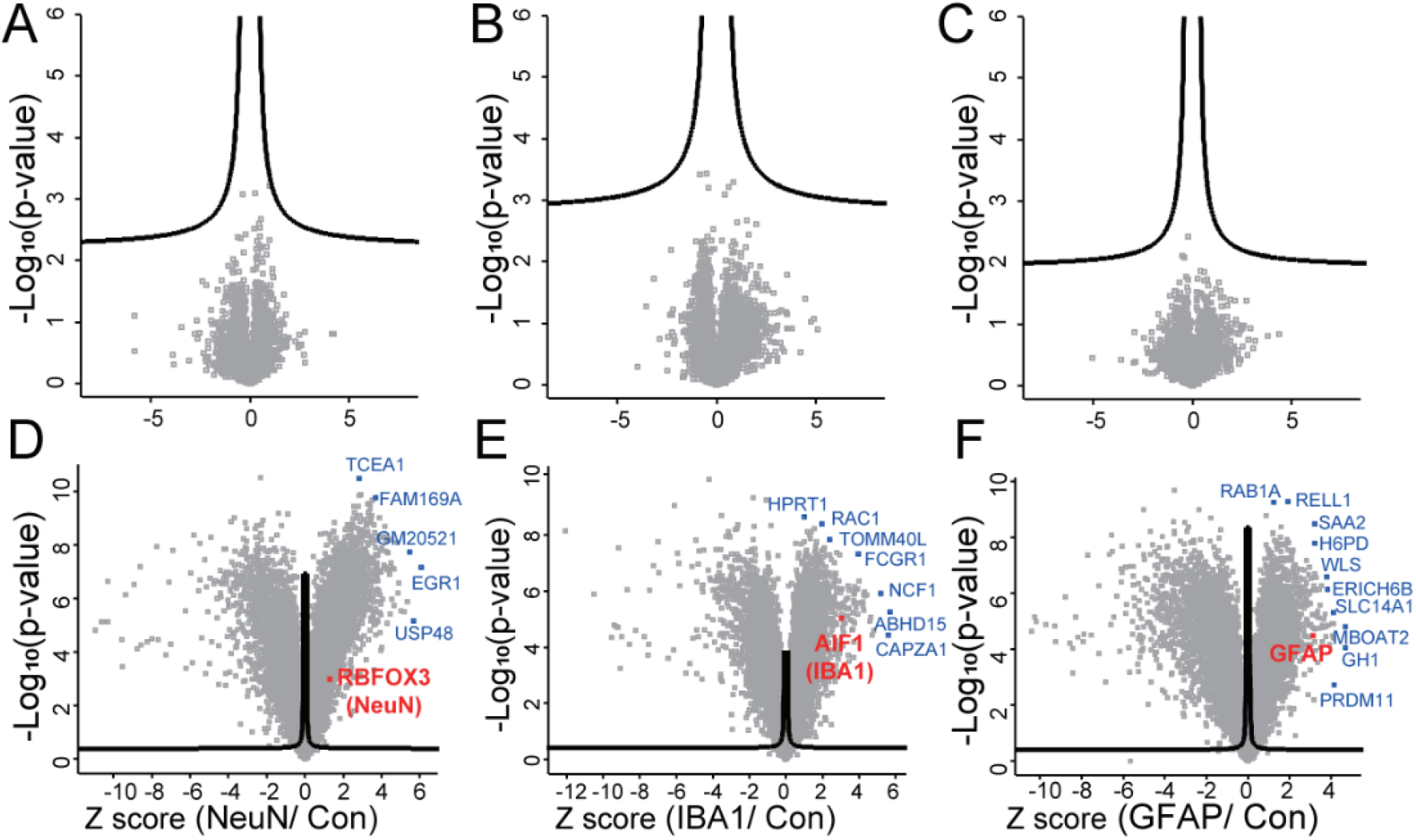
Volcano plot analysis results for iCAB proteomics data. The three volcano plots on the top (A-C) are of iCAB proteome analysis results for non-enriched samples. The three volcano plots on the bottom (D-F) are of iCAB proteome analysis results for enriched samples. The iCAB proteome data of anti-NeuN-antibody (NeuN; A and D), anti-IBA1 antibody (IBA1; B and E) and anti-GFAP antibody (GFAP; C and F) were compared to that of control (no primary antibody). The proteins outside the curved lines have *q*-values < 0.05.

**Supplementary Figure S6.**
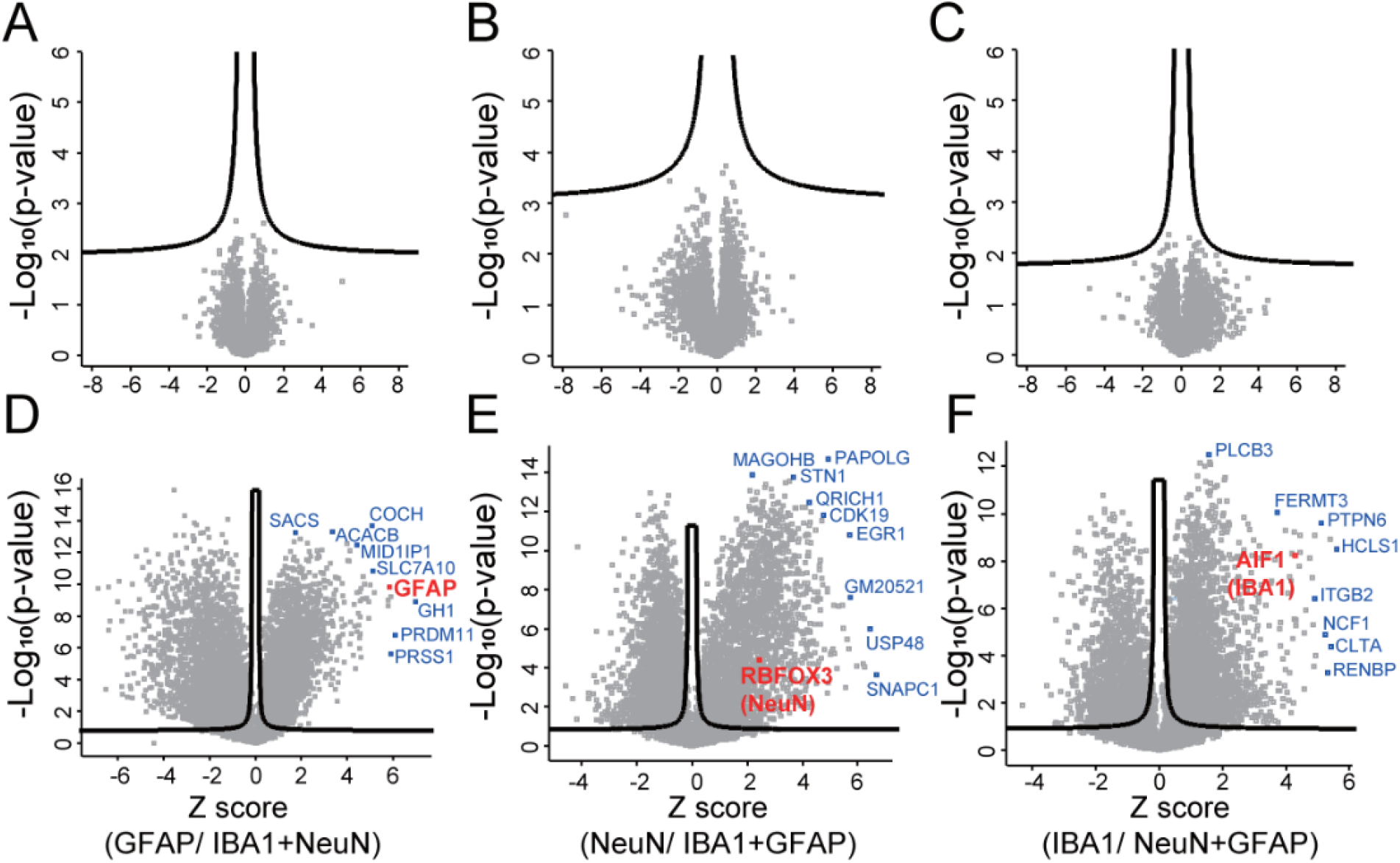
Volcano plot analysis results for iCAB proteomics data. The three volcano plots on the top (A-C) are of iCAB proteome analysis results for non-enriched samples. The three volcano plots on the bottom (D-F) are of iCAB proteome analysis results for enriched samples. The iCAB proteome data of anti-GFAP-antibody (GFAP; A and D), anti-NeuN antibody (NeuN; B and E) and anti-IBA1 antibody (IBA1; C and F) were compared to that of the other two cell types. The proteins outside the curved lines have *q*-values < 0.05.

## REFERENCES

1. Aebersold R, Mann M. Mass-spectrometric exploration of proteome structure and function. Nature. 2016;537(7620):347–55. Epub 2016/09/16. doi: 10.1038/nature19949. PubMed PMID: 27629641.

2. Gebreyesus ST, Siyal AA, Kitata RB, Chen ES, Enkhbayar B, Angata T, Lin KI, Chen YJ, Tu HL. Streamlined single-cell proteomics by an integrated microfluidic chip and data-independent acquisition mass spectrometry. Nat Commun. 2022;13(1):37. Epub 2022/01/12. doi: 10.1038/s41467-021-27778-4. PubMed PMID: 35013269; PMCID: PMC8748772.

3. Gstaiger M, Aebersold R. Applying mass spectrometry-based proteomics to genetics, genomics and network biology. Nat Rev Genet. 2009;10(9):617–27. Epub 2009/08/19. doi: 10.1038/nrg2633. PubMed PMID: 19687803.

4. Yang L, George J, Wang J. Deep Profiling of Cellular Heterogeneity by Emerging Single-Cell Proteomic Technologies. Proteomics. 2020;20(13):e1900226. Epub 2019/11/16. doi: 10.1002/pmic.201900226. PubMed PMID: 31729152; PMCID: PMC7225074.

5. Alvarez-Castelao B, Schanzenbacher CT, Hanus C, Glock C, Tom Dieck S, Dorrbaum AR, Bartnik I, Nassim-Assir B, Ciirdaeva E, Mueller A, Dieterich DC, Tirrell DA, Langer JD, Schuman EM. Cell-type-specific metabolic labeling of nascent proteomes in vivo. Nat Biotechnol. 2017;35(12):1196–201. Epub 2017/11/07. doi: 10.1038/nbt.4016. PubMed PMID: 29106408.

6. Hobson BD, Choi SJ, Mosharov EV, Soni RK, Sulzer D, Sims PA. Subcellular proteomics of dopamine neurons in the mouse brain. Elife. 2022;11. Epub 2022/02/01. doi: 10.7554/eLife.70921. PubMed PMID: 35098924; PMCID: PMC8860448.

7. Shuster SA, Li J, Chon U, Sinantha-Hu MC, Luginbuhl DJ, Udeshi ND, Carey DK, Takeo YH, Xie Q, Xu C, Mani DR, Han S, Ting AY, Carr SA, Luo L. In situ cell-type-specific cell-surface proteomic profiling in mice. Neuron. 2022;110(23):3882–96 e9. Epub 2022/10/12. doi: 10.1016/j.neuron.2022.09.025. PubMed PMID: 36220098; PMCID: PMC9742329.

8. Rayaprolu S, Bitarafan S, Santiago JV, Betarbet R, Sunna S, Cheng L, Xiao H, Nelson RS, Kumar P, Bagchi P, Duong DM, Goettemoeller AM, Olah VJ, Rowan M, Levey AI, Wood LB, Seyfried NT, Rangaraju S. Cell type-specific biotin labeling in vivo resolves regional neuronal and astrocyte proteomic differences in mouse brain. Nat Commun. 2022;13(1):2927. Epub 2022/05/26. doi: 10.1038/s41467-022-30623-x. PubMed PMID: 35614064; PMCID: PMC9132937.

9. MacDonald ML, Favo D, Garver M, Sun Z, Arion D, Ding Y, Yates N, Sweet RA, Lewis DA. Laser capture microdissection-targeted mass spectrometry: a method for multiplexed protein quantification within individual layers of the cerebral cortex. Neuropsychopharmacology. 2019;44(4):743–8. Epub 2018/11/06. doi: 10.1038/s41386-018-0260-0. PubMed PMID: 30390066; PMCID: PMC6372704.

10. Davis S, Scott C, Ansorge O, Fischer R. Development of a Sensitive, Scalable Method for Spatial, Cell-Type-Resolved Proteomics of the Human Brain. J Proteome Res. 2019;18(4):1787–95. Epub 2019/02/16. doi: 10.1021/acs.jproteome.8b00981. PubMed PMID: 30768908; PMCID: PMC6456870.

11. Dammer EB, Duong DM, Diner I, Gearing M, Feng Y, Lah JJ, Levey AI, Seyfried NT. Neuron enriched nuclear proteome isolated from human brain. J Proteome Res. 2013;12(7):3193–206. Epub 2013/06/19. doi: 10.1021/pr400246t. PubMed PMID: 23768213; PMCID: PMC3734798.

12. Hu P, Zhang W, Xin H, Deng G. Single Cell Isolation and Analysis. Front Cell Dev Biol. 2016;4:116. Epub 2016/11/09. doi: 10.3389/fcell.2016.00116. PubMed PMID: 27826548; PMCID: PMC5078503.

13. Bar DZ, Atkatsh K, Tavarez U, Erdos MR, Gruenbaum Y, Collins FS. Biotinylation by antibody recognition-a method for proximity labeling. Nat Methods. 2018;15(2):127–33. Epub 2017/12/20. doi: 10.1038/nmeth.4533. PubMed PMID: 29256494; PMCID: PMC5790613.

14. Killinger BA, Marshall LL, Chatterjee D, Chu Y, Bras J, Guerreiro R, Kordower JH. In situ proximity labeling identifies Lewy pathology molecular interactions in the human brain. Proc Natl Acad Sci U S A. 2022;119(5). Epub 2022/01/28. doi: 10.1073/pnas.2114405119. PubMed PMID: 35082147; PMCID: PMC8812572.

15. Rasband MN, Peles E, Trimmer JS, Levinson SR, Lux SE, Shrager P. Dependence of nodal sodium channel clustering on paranodal axoglial contact in the developing CNS. J Neurosci. 1999;19(17):7516–28. Epub 1999/08/25. doi: 10.1523/JNEUROSCI.19-17-07516.1999. PubMed PMID: 10460258; PMCID: PMC6782503.

16. Jang Y, Thuraisamy T, Redding-Ochoa J, Pletnikova O, Troncoso JC, Rosenthal LS, Dawson TM, Pantelyat AY, Na CH. Mass spectrometry-based proteomics analysis of human globus pallidus from progressive supranuclear palsy patients discovers multiple disease pathways. Clin Transl Med. 2022;12(11):e1076. Epub 2022/11/11. doi: 10.1002/ctm2.1076. PubMed PMID: 36354133; PMCID: PMC9647849.

17. Jang Y, Pletnikova O, Troncoso JC, Pantelyat AY, Dawson TM, Rosenthal LS, Na CH. Mass Spectrometry-Based Proteomics Analysis of Human Substantia Nigra From Parkinson’s Disease Patients Identifies Multiple Pathways Potentially Involved in the Disease. Mol Cell Proteomics. 2023;22(1):100452. Epub 2022/11/25. doi: 10.1016/j.mcpro.2022.100452. PubMed PMID: 36423813; PMCID: PMC9792365.

18. Kong AT, Leprevost FV, Avtonomov DM, Mellacheruvu D, Nesvizhskii AI. MSFragger: ultrafast and comprehensive peptide identification in mass spectrometry-based proteomics. Nat Methods. 2017;14(5):513–20. Epub 2017/04/11. doi: 10.1038/nmeth.4256. PubMed PMID: 28394336; PMCID: PMC5409104.

19. Ramachandran KV, Fu JM, Schaffer TB, Na CH, Delannoy M, Margolis SS. Activity-Dependent Degradation of the Nascentome by the Neuronal Membrane Proteasome. Mol Cell. 2018;71(1):169–77 e6. Epub 2018/07/07. doi: 10.1016/j.molcel.2018.06.013. PubMed PMID: 29979964; PMCID: PMC6070390.

20. Tyanova S, Temu T, Sinitcyn P, Carlson A, Hein MY, Geiger T, Mann M, Cox J. The Perseus computational platform for comprehensive analysis of (prote)omics data. Nat Methods. 2016;13(9):731–40. Epub 2016/06/28. doi: 10.1038/nmeth.3901. PubMed PMID: 27348712.

21. Tusher VG, Tibshirani R, Chu G. Significance analysis of microarrays applied to the ionizing radiation response. Proc Natl Acad Sci U S A. 2001;98(9):5116–21. Epub 2001/04/20. doi: 10.1073/pnas.091062498. PubMed PMID: 11309499; PMCID: PMC33173.

22. Kuleshov MV, Jones MR, Rouillard AD, Fernandez NF, Duan Q, Wang Z, Koplev S, Jenkins SL, Jagodnik KM, Lachmann A, McDermott MG, Monteiro CD, Gundersen GW, Ma’ayan A. Enrichr: a comprehensive gene set enrichment analysis web server 2016 update. Nucleic Acids Res. 2016;44(W1):W90–7. Epub 2016/05/05. doi: 10.1093/nar/gkw377. PubMed PMID: 27141961; PMCID: PMC4987924.

23. Hao Y, Hao S, Andersen-Nissen E, Mauck WM, 3rd, Zheng S, Butler A, Lee MJ, Wilk AJ, Darby C, Zager M, Hoffman P, Stoeckius M, Papalexi E, Mimitou EP, Jain J, Srivastava A, Stuart T, Fleming LM, Yeung B, Rogers AJ, McElrath JM, Blish CA, Gottardo R, Smibert P, Satija R. Integrated analysis of multimodal single-cell data. Cell. 2021;184(13):3573–87 e29. Epub 2021/06/02. doi: 10.1016/j.cell.2021.04.048. PubMed PMID: 34062119; PMCID: PMC8238499.

24. Hu BC. The human body at cellular resolution: the NIH Human Biomolecular Atlas Program. Nature. 2019;574(7777):187–92. Epub 2019/10/11. doi: 10.1038/s41586-019-1629-x. PubMed PMID: 31597973; PMCID: PMC6800388 Filtircine, SensOmics, Qbio, January, Mirvie, Oralome and Proteus. He is also on the scientific advisory board (SAB) of Genapsys and Jupiter and on the advisory board of the National Institute of Diabetes and Digestive and Kidney Diseases (NIDDK). P.V.K. serves on the SAB to Celsius Therapeutics. A.R. is a member of the SAB of ThermoFisher Scientific and Syros Pharmaceuticals and a founder and an equity holder of Celsius Therapeutics. A.R. holds various patents and has patent filings in the areas of single-cell and spatial genomic technologies, and is a member of the advisory council of the National Human Genome Research Institute (NHGRI). C.K. is a co-founder of Ocean Genomics. P.Y. is a co-founder, paid consultant, director, and equity holder of Ultivue and NuProbe Global and holds several patent filings in the areas of single-cell and spatial genomic technologies. N.G. is a co-founder and equity owner of Datavisyn. R.F.M. is a cofounder and board member of Quantitative Medicine and is on the Advisory Board of Predictive Oncology and the Scientific Advisory Board of the Morgridge Institute for Research. The other authors declare no competing interests.

25. Pang Z, Zhou G, Ewald J, Chang L, Hacariz O, Basu N, Xia J. Using MetaboAnalyst 5.0 for LC-HRMS spectra processing, multi-omics integration and covariate adjustment of global metabolomics data. Nat Protoc. 2022;17(8):1735–61. Epub 2022/06/18. doi: 10.1038/s41596-022-00710-w. PubMed PMID: 35715522.

26. Huang da W, Sherman BT, Lempicki RA. Systematic and integrative analysis of large gene lists using DAVID bioinformatics resources. Nat Protoc. 2009;4(1):44–57. Epub 2009/01/10. doi: 10.1038/nprot.2008.211. PubMed PMID: 19131956.

27. Allen NJ, Eroglu C. Cell Biology of Astrocyte-Synapse Interactions. Neuron. 2017;96(3):697–708. Epub 2017/11/03. doi: 10.1016/j.neuron.2017.09.056. PubMed PMID: 29096081; PMCID: PMC5687890.

28. Chen D, Shen X, Sun L. Strong cation exchange-reversed phase liquid chromatography-capillary zone electrophoresis-tandem mass spectrometry platform with high peak capacity for deep bottom-up proteomics. Anal Chim Acta. 2018;1012:1–9. Epub 2018/02/25. doi: 10.1016/j.aca.2018.01.037. PubMed PMID: 29475469; PMCID: PMC5831384.

